# HSV-1 Infection Alters *MAPT* Splicing and Promotes Tau Pathology in Neural Models of Alzheimer’s Disease

**DOI:** 10.1101/2024.10.16.618683

**Authors:** Emmanuel C Ijezie, Michael J Miller, Celine Hardy, Ava R. Jarvis, Timothy F Czajka, Lianna D’Brant, Natasha Rugenstein, Adam Waickman, Eain Murphy, David C Butler

**Author notes:** Correspondence Eain Murphy Ph.D. David Butler Ph.D.

## Abstract

**INTRODUCTION:** Herpes simplex virus 1 (HSV-1) infection alters critical markers of Alzheimer’s Disease (AD) in neurons. One key marker of AD is the hyperphosphorylation of Tau, accompanied by altered levels of Tau isoforms. However, an imbalance in these Tau splice variants, specifically resulting from altered 3R to 4R *MAPT* splicing of exon 10, has yet to be directly associated with HSV-1 infection

**METHODS:** To this end, we infected 2D and 3D human neural models with HSV-1 and monitored *MAPT* splicing and Tau phosphorylation. Further, we transduced SH-SY5Y-neurons with HSV-1 ICP27 which alters RNA splicing to analyze if ICP27 alone is sufficient to induce altered *MAPT* exon 10 splicing.

**RESULTS:** We show that HSV-1 infection induces altered splicing of *MAPT* exon 10, increasing 4R-Tau protein levels, Tau hyperphosphorylation, and Tau oligomerization.

**DISCUSSION:** Our experiments reveal a novel link between HSV-1 infection and the development of cytopathic phenotypes linked with AD progression.

**HIGHLIGHTS:**

- HSV-1 infection in forebrain organoids reduces the neurite length of MAP2-positive neurons.
- HSV-1 infection increases Tau hyperphosphorylation in both two-month-old and four-month-old forebrain organoids.
- HSV-1 infection increases Exon 10 containing (4R) *MAPT* mRNA and 4R-Tau protein expression in both forebrain organoids and human SH-SY5Y-neurons.
- HSV-1 ICP27 is both necessary and sufficient to induce increased 4R *MAPT* mRNA and 4R-Tau protein expression in SH-SY5Y-neurons.
- HSV-1 infection increases Tau oligomerization in both forebrain organoids and SH-SY5Y-neurons.

## 1. BACKGROUND

Alzheimer’s Disease (AD) is a neurological disorder that is characterized by progressive neurodegeneration, and accumulation of amyloid-ß (Aβ) plaques and aggregated Tau proteins.^1^ AD is currently the leading cause of dementia in the world. The economic impact of AD progression on the United States healthcare system is projected to be in excess of $360 billion in 2024 and this number is expected to continue rising.^2^ Currently, there is no known cure for AD, hence the need for further research into this disorder.

AD dementia is distinct from most types of dementia due to the presence of insoluble extracellular Aβ plaques and intracellular neurofibrillary Tau tangles.^3^ Aβ plaque accumulation is induced by the abnormal cleavage of the amyloid precursor protein (APP), a key player in neural synapse formation and repair. Tau neurofibrillary tangles (NFTs) are a major hallmark of disease pathology in tauopathies like AD. Tau is a microtubule-associated protein that is essential in maintaining microtubule stability within neuronal axons.^4–6^ During AD, Tau becomes hyperphosphorylated, dissociates from microtubules, and accumulates to form insoluble aggregates. These insoluble NFT aggregates impair neuronal function which leads to subsequent neuronal cell death. Recent studies have shown that aberrant forms of Tau oligomers can self-propagate and spread within the brain, leading to Tau protein misfolding and NFT formation in neighboring neurons.^7,8^ This spread is thought to occur in a “prion-like” manner, which may explain the spread of progressive neural degeneration across multiple brain regions, leading to widespread neuronal dysfunction and cognitive decline during AD.^7^ Standard of care therapeutics against AD are targeted toward reducing Aβ plaques, however, Tau pathology is recognized as a key driver of AD progression and has been shown to correlate more closely with neurodegeneration and cognitive decline than Aβ plaques.^9^ As such additional research is needed to better understand the role of Tau NFTs in AD pathology, to enable development of new and improved therapeutics and drug targets that can inhibit AD progression.

Tau is expressed from the microtubule associated protein tau (*MAPT*) gene as six splice variants which are grouped into two categories, based on the exclusion or inclusion of exon 10, resulting in either three repeats (3R-Tau) or four repeats (4R-Tau) of microtubule-binding domains.^10,11^ In healthy adult humans, the stoichiometric ratio of 3R to 4R Tau is 1:1 and this ratio becomes altered in individuals diagnosed with AD.^12^ The balance in 3R:4R Tau protein expression is regulated by the alternative splicing machinery, which includes the Serine Arginine Splice Factor 1 (SRSF1) and its respective kinase Serine Arginine protein Kinase 1 (SRPK1), which influence the inclusion or removal of *MAPT* exon 10.^13^ The imbalance in the 3R:4R Tau ratio has been shown to induce increased phosphorylation and oligomerization of Tau.^10,14^ Further research is needed to fully elucidate potential factors that may be contributing to altered splicing of *MAPT* in AD patients.

Familial genetic mutations in genes *APP*, Presenilin 1 and 2, and Apolipoprotein E, confer an increased risk of developing AD.^15^ While these mutations increase the likelihood of developing AD, they do not directly cause the disease. There is growing interest in investigating etiological factors that may have a causal link to AD development. Recently, neurotrophic pathogens like bacteria and viruses have been linked to AD progression. A study in Taiwan showed that patients infected with herpesviruses had an increased risk of developing AD compared to uninfected patients.^16^ Importantly, individuals that were infected with herpesviruses and were undergoing antiviral therapy to limit viral replication and reactivation did not show an increased risk of AD thus suggesting a link between the virus and the disease.^16^ Another multiscale study performed on postmortem tissues from AD patients showed that herpes simplex virus 1 (HSV-1) viral DNA, RNA, and proteins could be detected in brain regions that had elevated markers for AD.^17^ These studies add to the growing evidence from murine models of AD showing that HSV-1 infection induces pathological hallmarks of AD progression.^18,19^

HSV-1 is a neurotropic herpesvirus that establishes a latent infection cycle in neurons. HSV-1 infection in mice is increasingly associated with Aβ accumulation and neuroinflammation, both major pathological features in AD.^19^ Evidence from murine models suggests that HSV-1 infection may contribute to amyloid-beta (Aβ) accumulation by disrupting pathways involved in APP processing via beta and gamma secretase.^20^ These studies strengthen the link between HSV-1 infection and the development of neurological diseases like AD. HSV-1 expresses an immediate early protein named Infected Cell Protein 27 (ICP27) that interacts with the host cell alternative splicing machinery to alter the splicing of cellular and viral transcripts.^21^ HSV-1 ICP27 protein interacts with SRPK1 through its RGG box.^22^ This interaction leads to decreased phosphorylation of SRSF1 leading to altered splicing of cellular RNA transcripts.^22^ Dysregulation of SRPK1 and SRSF1 interaction by ICP27 alters alternative splicing of the promyelocytic leukemia gene (PML), which is a key player in cell-intrinsic immunity.^23^ Interestingly, the regulation for exon 10 inclusion or exclusion during *MAPT* alternative splicing is also controlled by the SRSF proteins, and so this presents an opportunity for HSV-1 ICP27 protein to disrupt splicing of *MAPT*.

Animal models of AD such as the 5xFAD mice, effectively replicate the pathology associated with familial or genetically linked AD, representing only a small subset of AD.^24,25^ However, adult mice naturally express only three isoforms of 4R-Tau compared to humans who express six (three 3R and three 4R isoforms), thus are not suitable models to evaluate the role of alternative splicing of *MAPT* thereby leading to Tau pathology.^26^ To study potential causes of sporadic AD, which accounts for the majority of AD cases, it is crucial to have a model that is capable of recapitulating the physiological makeup of a healthy human brain. HSV-1 is a human virus with a close relationship with its host and as such infections in mice may not accurately reflect the pathogenesis of the virus in its natural host environment. Recently, brain organoids have emerged as a vital 3D *in vitro* system to study the role of viral infections and genetic mutations in causing neurological disease.^27,28^ These brain organoids model the 3D architecture of the human brain and have both glial and neural cell types, similar to what is found in humans.^27,29^ Brain organoids are utilized to model HSV-1 induced virus encephalopathy in humans and as such represent an ideal *in vitro* model for studying virus induced disease in the brain.^30^

To test if HSV-1 infection alters splicing of *MAPT* mRNA we infected iPSC-derived forebrain organoids and neurons and analyzed splicing of *MAPT* exon 10 and protein levels of 4R-Tau. We observed robust HSV-1 infection in both neurons and glial cells within our organoids. HSV-1 infected organoids have upregulated phosphorylated Tau and have an increased expression of both 4R *MAPT* mRNA over total *MAPT* mRNA expression, and increased 4R-Tau protein level. We next infected organoids with HSV-1 that lacked the immediate early protein ICP27 because of its role in disrupting alternative splicing machinery. We observed that in the absence of ICP27, the increase in altered splicing of *MAPT* was not observed coupled to 4R-Tau protein levels being similar to uninfected samples, indicating that ICP27 was necessary for this response. Further we transduced SH-SY5Y-neurons with ICP27 and observed a marked increase in the 4R-Tau protein levels indicating that the protein ICP27 was also sufficient in inducing this response. Finally, we also observed increased Tau oligomerization in both 2D-neurons and 3D organoids infected with HSV-1. Together our findings provide evidence of a common human pathogen negatively impacting a key component of AD progression.

## 2. METHODS

### 2.1. Cells and Viruses

Vero cells were grown in Dulbecco’s medium (Cat No. 11500P. Cleveland clinic Cell services Media) with 10% FBS. HSV-1 Patton strain with a GFP reporter, a kind gift from Ian Mohr,^31^ was expanded and passaged in Vero cells. HSV-1 F strain with a tdTomato reporter on an SV40 promoter with immediate early viral gene expression kinetics, a gift from Greg Smith was expanded and passaged in Vero cells also.^32^ The recombinant HSV-1 F strain with ICP27 deleted was made using the GALK recombineering protocol by Warming et al.,^33^ and expanded in complementary cell line Vero 27 cells, a kind gift from Dr. Stephen Rice.^34^

### 2.2. SH-SY5Y Cell Culture and Differentiation

SH-SY5Y cells were maintained and differentiated into neurons according to the protocol described by Shipley et al., with slight modifications.^35^ On Day 0, SH-SY5Y cells were passaged from a T75 flask. Basic media (EMEM, 15% heat-inactivated fetal bovine serum (FBS), and 1X Anti-Anti) and Trypsin were warmed in a 37°C bead bath. Cells were washed with 1X PBS, trypsinized (2 mL of 0.25% Trypsin-EDTA) and incubated at 37°C for 5 minutes. Trypsin was quenched by adding 1 mL of media per well, and the cells were resuspended by gentle trituration and collected in a sterile 50 mL conical tube. Cells were centrifuged (300 x g for 5 minutes), resuspended in Basic Media, and counted using Trypan Blue. Cells were plated at a density of 400,000 cells per well in a 6-well plate and incubated at 37°C with 5% CO2.

On Day 1, differentiation was initiated with Differentiation Media #1 (EMEM, 2.5% heat-inactivated FBS, 1X Anti-Anti, and 10 µM retinoic acid (RA) added immediately before feeding). The media was changed every 2 days. On Day 7, cells were trypsinized (500 µL of 0.05% trypsin, diluted from 0.25% trypsin with 1X CMF-PBS, added per well), incubated at 37°C for 5 minutes, quenched by adding 1 mL of media per well, resuspended by gentle trituration, and collected in a sterile 50 mL conical tube. Cells were centrifuged (300 x g for 5 minutes), resuspended in Differentiation Media #1 with RA, and replated. Plates were coated with PLO (0.01%) overnight and Laminin 2020 (1:100 dilution in 1X PBS+CM) at 4°C overnight.

On Day 8, the media was changed to Differentiation Media #2 (EMEM, 1% heat-inactivated FBS, 1X Anti-Anti, and 10 µM RA). Cells were fed every 2 days.

On Day 10, cells were trypsinized (500 µL of 0.05% trypsin, diluted from 0.25% trypsin with 1X PBS, added per well), incubated at room temperature for 5 minutes, quenched by adding 1 mL of media per well, resuspended by gentle trituration, and collected in a sterile 50 mL conical tube. Cells were centrifuged (300 x g for 5 minutes), resuspended in Differentiation Media #2 with RA, and replated. Plates were prepared as described earlier.

On Day 11, media was changed to Differentiation Media #3 (Neurobasal, 1X B-27, 20 mM KCl, 1X Anti-Anti, 2 mM GlutaMAX, 50 ng/mL BDNF, 2 mM db-c-AMP, and 10 µM RA added immediately before feeding). Media changes were performed every 2-3 days for another week until treatment.

### 2.3. ICP27 Plasmid Design and Lentivirus Production

ICP27 was synthesized by Vector Builder and cloned into an inducible tetracycline on 3rd generation lentiviral vector (pLV[Exp]-EGFP:T2A: Puro-TRE>{ICP27_Genbank AB235841.1}). The tetracycline-on system utilized in this study is dependent on the reverse tetracycline-controlled transactivator (pLV[Exp]-CMV>tTS/rtTA/Hygro (VectorBuilder: Vector ID: VB010000-9369xhm). Lentiviruses were produced in 293FT cells (ThermoFisher; Cat# R70007) following the protocol by D’Brant, Rugenstein et al.. ^36^

### 2.4. Generating 3D forebrain organoids derived from human iPSCs

Human iPSCs were cultured on Matrigel-coated 6-well plates with mTeSR medium following standard protocols. Organoids were produced by the NeuraCell Core facility using established methods.^37,38^ Briefly, iPSCs were expanded to 70-80% confluency, dissociated into single cells for spheroid formation, and resuspended with ROCK inhibitor. Cells were seeded into a 96-well plate and fed daily with E6 medium and growth factors before transitioning to Neurobasal A medium with reduced growth factors. On day 20, organoids underwent quality control for proper cortical forebrain patterning before transitioning to less frequent feedings. Organoids were cultured for up to either two or four months.

### 2.5. Virus Infections and Lentivirus Transduction

GFP-tagged HSV-1 was used to infect 2-month-old iPSC-derived 3D forebrain organoids for 48 hours. td Tomato-tagged HSV-1 was used to infect 4-month-old iPSC-derived 3D forebrain organoids for 48 hours at a multiplicity of infection (MOI) of 0.1 and SH-SY5Y differentiated neurons (SH-SY5Y-neurons) for 24 hours at an MOI of 1.

For lentiviral transduction, lentiviral vectors were utilized to transduce SH-SY5Y-neurons at a MOI of 1 for 24 hours. Subsequently doxycycline was added every 24 hours for 72 hours.

### 2.6. Organoid Processing for Immunofluorescence Analysis

Forebrain Organoids were fixed in 4% Paraformaldehyde in 1X PBS for 30 minutes at 37 C. After fixation, organoids were suspended in sucrose and subsequently frozen in OCT. Frozen organoid blocks were sectioned on a cryostat at 15 um thickness for all sections.

### 2.7. Immunofluorescence Analysis

Forebrain organoid sections and 2D SH-SY5Y-neurons were incubated overnight at 4°C with primary antibodies in 2.5% Bovine Serum Albumin (BSA) and 0.05% Triton X in 1X PBS (See Table 1), washed with 1X PBS and followed by secondary antibody staining for 1 hour and imaging on a Nikon Olympus microscope. Immunofluorescence signal was quantified using the NIH Image J software. An average of 6 fields of view were quantified for each protein analyzed.

**Table 1.** List of Primary and Secondary antibodies

|  |  |  |  |  |  |
| --- | --- | --- | --- | --- | --- |
| WESTERN BLOT |  |  | WESTERN BLOT |  |  |
| Primary AB | Catalogue Number | Supplier | Secondary AB | Catalogue Number | Supplier |
| ICP27 | P1113 | Virusys | Anti-Mouse IgG, HRP-linked Antibody | 31430 | Cell Signalling Technology |
| Tau 4R | 30328S | Cell Signalling Technology | Anti-Rabbit IgG, HRP-linked Antibody | 7074S | Cell Signalling Technology |
| Total Tau | MA5-41098 | Thermofisher |  |  |  |
| VP16 | sc-7545 | Santa Cruz |  |  |  |
| B-Actin | 4967S | Cell Signalling Technology |  |  |  |
| IFA |  |  | IFA |  |  |
| Primary AB | Catalogue Number | Supplier | Secondary AB | Catalogue Number | Supplier |
| Tau | MA5-41098 | Thermofisher | Anti-mouse IgG (H+L), F(ab')2 Fragment (Alexa Fluor® 488 Conjugate) | 4408S | Cell Signalling Technology |
| AT8 | MN1020 | Thermofisher | Anti-rabbit IgG (H+L), F(ab')2 Fragment (Alexa Fluor® 488 Conjugate) | 4412S | Cell Signalling Technology |
| Tau 4R | ET3 | Peter Davies |  |  |  |
| PThr231 | RZ3 | Peter Davies |  |  |  |
| MAP2 | 13-1500 | Thermofisher |  |  |  |
| NEUN | MAB377 | Millipore Sigma |  |  |  |
| Nestin | MAB1259 | R&D |  |  |  |
| GFAP | GFAP | Aves Lab |  |  |  |
| T22 | ABN454 | Millipore Sigma |  |  |  |
| b-Tubulin III | MA1-118 | Thermofisher |  |  |  |

### 2.8. Organoid Size Measurement

Bright-field images of four forebrain organoids infected with either Mock or HSV-1 were used to measure the average diameter of organoids using the NIH Image J software.

### 2.9. Neurite Length Measurement

Forebrain organoid sections were stained with the neuron marker microtubule-associated protein 2 (MAP2) to evaluate neural cell morphology. Average neurite length was measured using NIH Image J software. 9 images from three biological replicates were analyzed and the average neurite length was calculated for each image. 9 average neurite lengths were then graphed on the GraphPad prism program with the mean result indicated.

### 2.10. Western Blot Protein Analysis

Whole organoids were lysed in RIPA buffer and mixed with sample loading buffer. Samples were then boiled for 5 minutes and then spun down and equivalent amount of protein was loaded onto 10% SDS PAGE gels. Membranes were probed for primary antibodies overnight at 4 degrees Celsius. Blots were washed three times (5 minutes each) with 0.1% Tween in 1X Tris-buffered saline buffer, and the secondary antibody was added for an hour. After incubation, blots were washed and developed with a chemiluminescence detection kit. See Table 1 for all antibody information.

### 2.11. Quantitative Reverse-Transcription-Polymerase Chain Reaction

Total RNA was isolated from whole forebrain Organoids (average of 3 individual organoids), and SH-SY5Y-neurons with Trizol (Invitrogen) RNA isolation reagent per the manufacturer’s instructions. The RNA product was precipitated using Isopropanol and 0.5-1ug of RNA was then reverse transcribed using the TaqMan RNA to cDNA kit (Applied Biosciences). The reverse transcription cDNA product was then analyzed using reverse transcription polymerase chain reaction (RT-PCR) TaqMan gene expression assay probes, on a CFX Connect QPCR machine (Bio-rad). The TaqMan gene expression probes utilized were: 4R *MAPT* Exon 10 (Hs00902312_m1), Total *MAPT* (Hs00902194_m1), and hypoxanthine phosphoribosyl transferase 1 (*HPRT1*, Reference gene, Hs02800695_m1) mRNA. 4R *MAPT* mRNA expression was calculated by analyzing the percentage of total *MAPT* mRNA that contains Exon 10.

### 2.12. SCRNA Sequencing

Organoid Dissociation for 10X processing: Organoids were dissociated using the Papain Dissociation System kit (Worthington) following the manufacturer’s instructions. Briefly, organoids were incubated in a 5.0 mL conical tube (Eppendorf) containing 750 µL of papain and Earle’s Balanced Salt Solution (EBSS) supplemented with Deoxyribonuclease I (DNase) at 37°C with gentle shaking (150 rpm). The tubes were shaken at 37°C and 150 rpm for 15 minutes, after which the organoids were triturated using wide-bore tips. After an additional 15 minutes of digestion, the organoids were triturated again with shaking at 37°C and 150 rpm. This digestion and trituration cycle was repeated two more times. Afterward, 500 μL of EBSS, 30 uL of DNase I solution, and 150 μL of pre-prepared Worthington inhibitor solution were added, and the mixture was gently mixed. The cells were centrifuged at 400 x g for 10 minutes at room temperature, rinsed once with 1X PBS to remove debris, and resuspended in 1X PBS containing 0.1% BSA to a cell concentration. Cells were manually counted with a hemocytometer and resuspended to a concentration of 1 x 106 cells/ml. The cell suspension was stored on ice until 10X processing. For the RNA Integrity Number (RIN) determination, 100ul of cells were removed and centrifuged at 400 x g for 10min at 37°C. Pellets were then snap-frozen and stored at -80°C until use.

Single cell RNA sequencing: Single cell RNA sequencing analysis was performed using the 10x Genomics Next GEM Single Cell 3’ Reagent kits according to the manufacturer’s recommended protocol. Sample quality and library quantity was assessed using an Agilent 4200 TapeStation with High Sensitivity D500 ScreenTape Assay and Qubit Fluorometer (Thermo Fisher Scientific) prior to sequencing anaylsis. Sequencing was performed on an Illumina NextSeq 2000 instrument (Illumina) using P3 reagent kits (100 cycles) set at the following sequencing parameters: 28 cycles for Read1, 10 cycles for Index1, 10 cycles for Index2, and 90 cycles for Read2.

5’ gene expression analysis and visualization: Sample demultiplexing, alignment, and barcode/UMI filtering was performed using the Cellranger software package v5.0 (10x Genomics, CA) using the mkfastq and count commands, respectively. Gene alignment was performed using a custom reference genome created using the Cellranger mkref implementation, combining the sequence for tdTomato and the HSV-1 genome (NC_001806.2) as an additional chromosome to the human reference genome (Ensembl GRCh38.93). Gene expression analysis and visualization was performed in R studio (v4.3.2) using R package Seurat (v4.4.0). Sample datasets were filtered to contain cells with less than 20% mitochondrial RNA content and between 750-5000 unique features. Genes expressed in fewer than 3 cells were excluded from analyses. Cell cycle scores were calculated using CellCycleScoring(), and the percentage of transcripts mapped to HSV-1 or tdTomato were computed for all cells. Data scaling, normalization and transformation were performed using the Seurat SCTransform() function, where cell cycle, HSV-1 expression, and tdTomato expression were regressed out. Following this, conserved features in the dataset were identified using SelectIntegrationFeatures() and PrepSCTIntegration(). Multi-sample integration was then performed using the FindIntegrationAnchors() and IntegrateData() functions, with the Mock sample used as the reference dataset. Following data normalization, principal component analysis was performed using RunPCA(). Cells were then clustered using default parameters, based on the first 10 principal components and with a resolution parameter of 0.1 using FindNeighbors() and FindClusters(). Automated cell type identification was performed using the R-package ScType.^39^ A custom template for annotation was created using the ScType database cell-specific marker expression of expected cell populations in brain tissue, with the addition of select genes according to previously published characterization of the organoid model used.^40^

### 2.13. Statistical Analysis

Statistical tests were performed on all figures using GraphPad Prism 10. P-values and their respective statistical tests are indicated in each figure legend.

## 3. RESULTS

### 3.1. HSV-1 Predominantly infects Astrocytes, Neurons, and Neuroepithelial Cells in Human 3D Forebrain Organoids

Brain organoids model the developing human brain and provide a useful tool for studying neurological diseases. To better understand the cellular tropism and neuropathological effects of HSV-1 infection in AD-relevant cell types, we utilized 3D forebrain organoids derived from human iPSCs to model HSV-1 infection in a human brain. We cultured human iPSC-derived 3D organoids for two months and proceeded to infect them for 48 hours with a fluorescently tagged HSV-1 **(Figure 1A).** After 48 hours of infection, we observed widespread green fluorescent protein expression (GFP) in organoids infected with HSV-1, but did not observe fluorescence in the Mock infected organoids **(Figure 1B).** We next measured the average diameter of our two groups (Mock and HSV-1 infected organoids) to determine if HSV-1 infection induced gross necrosis. Comparing the average diameter, we observed no significant difference between the two groups **(Figure 1C).** This indicates that HSV-1 infected organoids are still intact and not necrotic despite infection allowing for optimal analysis of virus induced pathology. Next, we sectioned HSV-1 infected organoids to investigate the cellular tropism of our GFP-tagged HSV-1. Immunofluorescence analysis using neural cell marker neuronal nuclear antigen (NeuN), glial cell marker glial fibrillary associated protein (GFAP), and neural progenitor cell marker nestin revealed expression of these three markers respectively in cells positive for the viral GFP signal. **(Figure 1D).** This result indicates that HSV-1 can infect neural progenitor cells, glial cells, and neurons within forebrain organoids, all known sites of natural infection.

**Figure 1.**
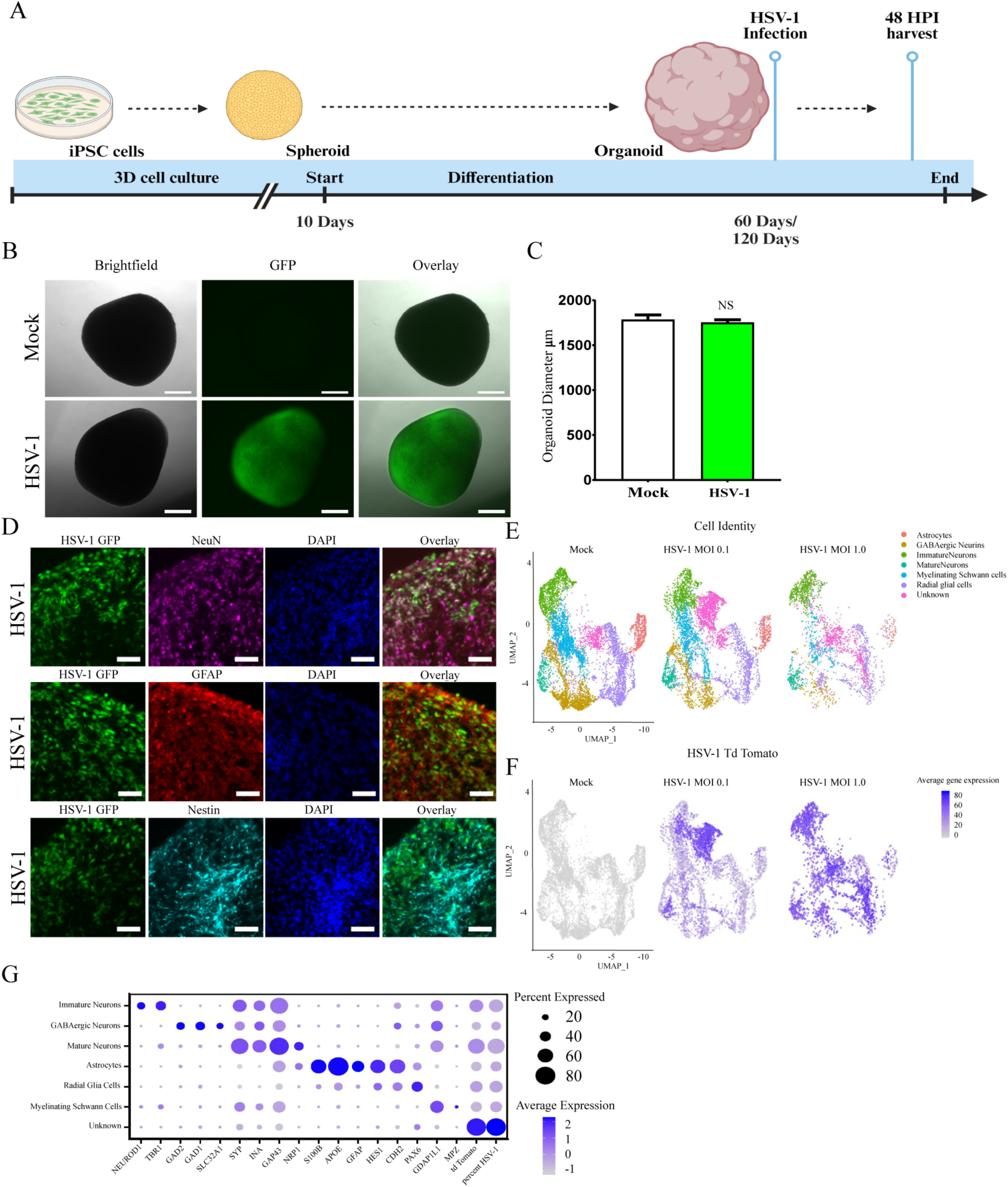
HSV-1 infects Astrocytes, Neurons and Neuroepithelial cells in Forebrain Organoids. A. Schematic diagram of 3D forebrain organoid culture with timeline for HSV-1 infection at 2-month-old or 4-month-old. B. 2-month-old organoids were either infected with Mock or infected with HSV-1 expressing the *eGFP* gene at an MOI of 0.5. HSV-1 eGFP expression at 48 hours post infection (hpi) in brain organoids was analyzed by epifluorescence imaging. Scale bar 500 μm. C. The average organoid diameter between treatment groups was measured and plotted. N of 3. Unpaired Student t test. D. Immunofluorescence analysis of neural cell marker (NeuN), glial cell marker (GFAP) and neural precursor cell marker (Nestin) in HSV-1 infected organoids. GFP (HSV-1). Scale bar 50 μm. E. Integrated uniform manifold approximation and projection (UMAP) of of scRNA seq data indicating annotated cell types in both Mock and tdTomato tagged HSV-1-infected 4-month-old organoids. F. Feature plot of tdTomato mRNA transcript in Mock and HSV-1-infected organoids. G. Dot plot indicating expression of key markers for the annotation and of cell types identified in scRNA seq of organoids. NS = P. Value > 0.05. P* ≤ 0.05, P** ≤ 0.01, P*** ≤ 0.001, P**** ≤ 0.0001. Error bars are the standard error of the means (SEM), Multiplicity of Infection (MOI).

Next, in order to investigate cellular tropism of HSV-1 in a model reflecting a more advanced stage of brain development, we generated older organoids containing further differentiated cell types such as mature neurons and astrocytes. We cultured forebrain organoids for 4 months and then infected them with HSV-1 encoding a tdTomato fluorescent reporter cassette driven by an immediate early promoter at an MOI of 0.1 or 1 for 48 hours and subsequently performed single-cell RNA sequencing on both the HSV-1 infected and Mock-infected organoids. The tdTomato tagged HSV-1 allows for identification of all infected cells that have undergone immediate early gene expression, including cells that have had abortive infections. Uniform manifold approximation and projection (UMAP) of scRNAseq data and cell type annotations based on the expression of key lineage defining markers identified the presence of multiple neuronal-linage cell types within both the HSV-1 and Mock-infected 3D forebrain organoids, including neurons, astrocytes and neural progenitor cells (Radial glia) **(Figure 1E and G).** Consistent with the infection conditions utilized in this experiment, HSV-1 and tdTomato transcripts were detected in increasing-abundance in the organoids infected at an MOI of 0.1 and 1, respectively, while no viral transcripts were identified in the Mock-infected sample **(Figure 1F)**. HSV-1 and tdTomato transcripts were observed in all annotated cell types in the organoids to varying degrees with the highest expression found in neurons and astrocytes as predicted. These results suggest that in this iPSC-model system multiple cell types: including neurons, astrocytes, and neural progenitor cells, are both susceptible and permissive to HSV-1 infection **(Figure 1E**, **F and G)**. These data collaborate with what we observed in the sectioned two-month-old organoids and concludes that HSV-1 infects both neural and glial cell types in forebrain organoids.

### 3.2. HSV-1 Infection Induces Neuropathology in Forebrain Organoids

HSV-1 has previously been shown to induce neuropathology, including Aβ accumulation and neuroinflammation in unpatterned brain organoids.^41,42^ In this study, we aimed to investigate whether HSV-1 infection also induced neuropathology to neuronal processes, potentially leading to loss of neural connections and synapses, a key phenotype of AD pathology. We performed immunofluorescence analysis of the neuronal marker microtubule-associated protein 2 (MAP2), which is primarily localized in dendrites, on Mock and HSV-1 infected 4-month-old forebrain organoids. Our initial analysis indicated a phenotypic difference in neurite outgrowth in our Mock versus HSV-1 infected organoids **(Figure 2A)**. Neurons in HSV-1 infected organoids displayed aberrant neurite morphology, with most MAP2-positive neurites being shortened and irregular in shape when compared to Mock infected controls **(see white arrow in Figure 2A).** Using NIH Image J software, we outlined and measured the average neurite length in both Mock and HSV-1 infected organoid sections. Quantification of the outlined neurites revealed that HSV-1 infected neurons had a significant reduction in neurite length, likely due to neurite retraction, a major phenotype of neural pathology **(Figure 2B and C).** In conclusion, our findings demonstrate that HSV-1 infection induces neuropathology which impairs the maintenance of neuronal processes in organoids

**Figure 2.**
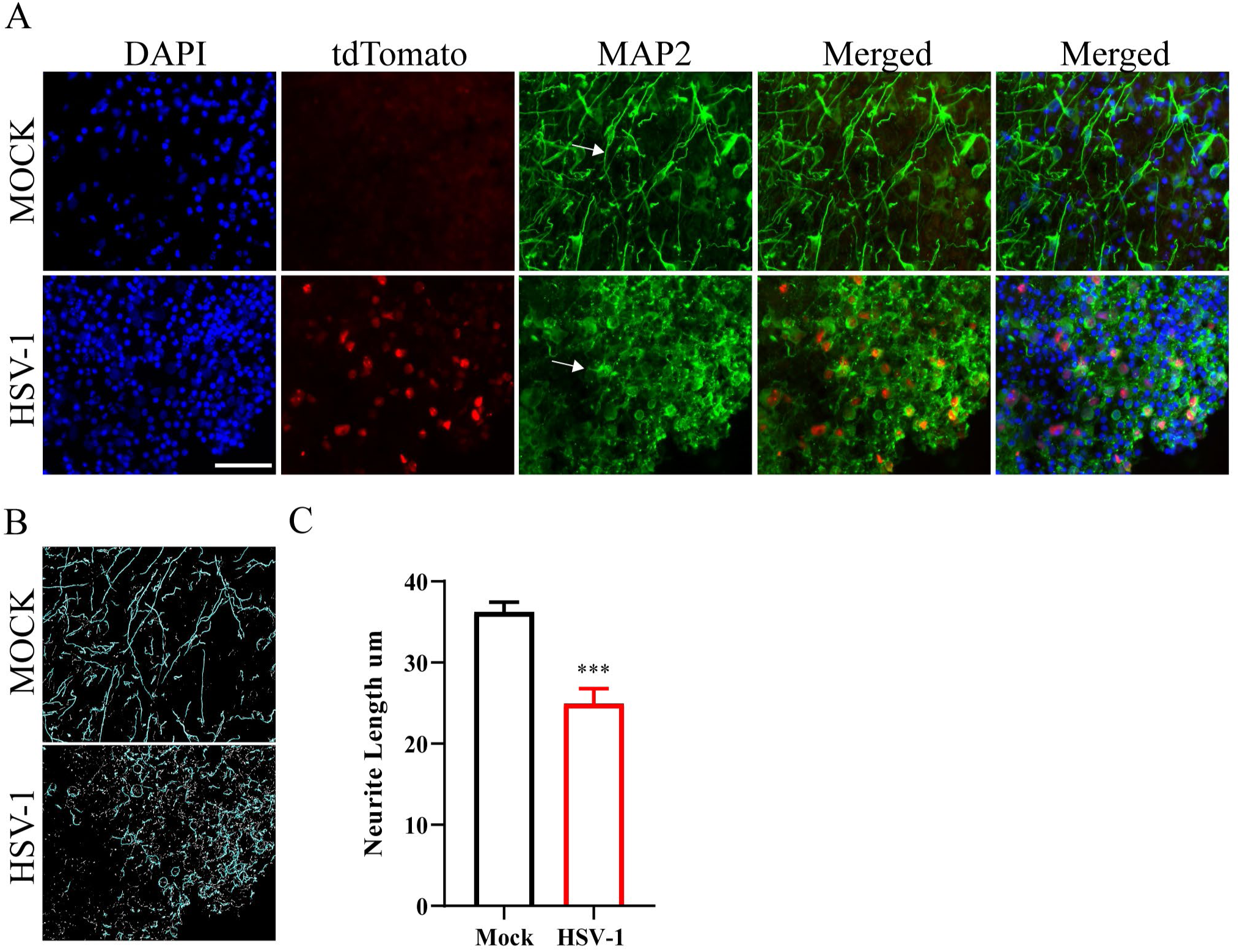
HSV-1 Infection Reduces Neurite Length in Forebrain Organoids. **A**. Immunofluorescence analysis of MAP2 staining in Mock and HSV-1 infected organoids. B. Image J neurite outline from both Mock and HSV-1 infected organoids. C. Average neurite length quantification of MAP2 positive neurons in Mock and HSV-1 infected organoids. N of 3, unpaired Student t test NS = P. Value > 0.05. P* ≤ 0.05, P** ≤ 0.01, P*** ≤ 0.001, P**** ≤ 0.0001. Error bars are SEM. Scale bar is 50 um.

### 3.3. HSV-1 Infection Induces Increased Phosphorylation of Tau in Forebrain Organoids

HSV-1 infection induces Tau hyperphosphorylation in 2D neurons.^43,44^ Therefore, we analyzed phosphorylation of Tau at residues Serine 202, Threonine 205, and Threonine 231 in our HSV-1 infected organoids. Using the phosphorylated Tau AT8 antibody that recognizes phosphorylated residues at Serine 202 and Threonine 205 and the phosphorylated Tau antibody RZ3 that recognizes phosphorylated residues at Threonine 231, we performed immunofluorescence analysis on 2-month-old and 4-month-old organoid sections respectively. The immunofluorescence analysis revealed a statistically significant increase in AT8 staining in HSV-1 infected 2-month-old organoids compared to Mock-infected controls, indicating an upregulation of Tau phosphorylation in response to HSV-1 infection of organoids **(Figure 3A and B).** Additionally, we stained the same sections for total Tau to observe changes to total Tau protein within infected organoids. We also observed a statistically significant increase in total Tau staining in the HSV-1 infected organoids **(Figure 3A and C).** Furthermore, our immunofluorescence analysis of 4-month-old organoids showed a significant increase in phosphorylated Threonine 231 staining in HSV-1 infected sections **(Figure 3D and E).** These results demonstrate that HSV-1 infection drives increased Tau phosphorylation, a process implicated in the development of neurodegenerative diseases.

**Figure 3.**
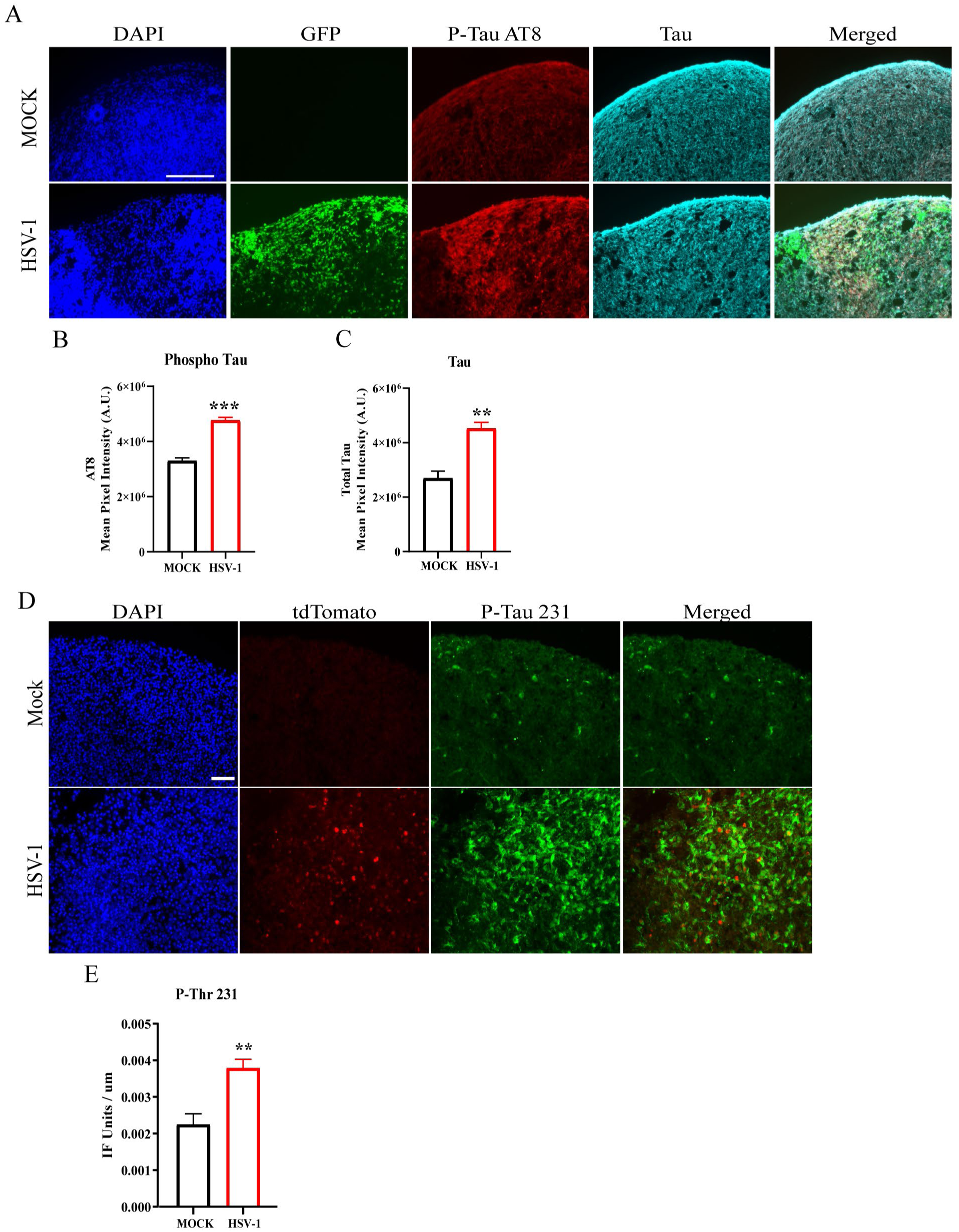
HSV-1 Infection Induces Tau Hyperphosphorylation in Forebrain Organoids. A. Immunofluorescence analysis of Tau and phosphorylated Tau AT8 on two-month-old organoid sections infected with either Mock or HSV-1 tagged with GFP. B and C. Quantification of immunofluorescence signal from A. N of 3. D. Immunofluorescence analysis of phosphorylated-Tau Threonine 231 staining in both Mock and HSV-1 (tdTomato tagged) infected organoids. E. Quantification of immunofluorescence signal from D. N of 3. Statistical analysis for graphs, used unpaired Student t test NS = P. Value > 0.05. P* ≤ 0.05, P** ≤ 0.01, P*** ≤ 0.001, P**** ≤ 0.0001. Error bars are SEM. Scale bar 50 um.

### 3.4. HSV-1 Infection Induces Increased 4R-Tau Expression in Forebrain Organoids

Increased expression of 4R-Tau in humanized mice, designed to contain both human 3R and 4R Tau isoforms, induces elevated hyperphosphorylation of Tau.^45^ Thus, we investigated if HSV-1 infection could increase 4R *MAPT* mRNA expression in our human derived 3D forebrain organoids. We hypothesized that this increase may be due to the impact of HSV-1 altering splicing of *MAPT* thereby increasing the percentage of total *MAPT* mRNA that would be 4R *MAPT* mRNA and an increase in 4R-Tau protein isoform pathology. To determine if HSV-1 alters splicing of *MAPT* mRNA of infected organoids, we utilized TaqMan probes specific for either total *MAPT* mRNA or *MAPT* exon 10, the spliced mRNA isoform that is found only in 4R *MAPT* mRNA. We performed RT-PCR on both Mock and HSV-1 infected 2 and 4-month-old organoids harvested at 48 hours post infection (hpi). Our RT-PCR results show that total *MAPT* mRNA from 2-month-old HSV-1 infected organoids was significantly reduced when compared to Mock **(Figure 4A)**. This finding was expected as host mRNA transcripts are reduced in levels by HSV-1 virion host shutoff protein (VHS) which functions in degrading host mRNA transcripts early during the virus life cycle.^46^ Importantly, analysis of exon 10 containing 4R *MAPT* mRNA revealed that HSV-1 infected organoids have a threefold increase in the percentage of *MAPT* mRNA that contained exon 10 (4R *MAPT*) when compared to Mock **(Figure 4B)**. Furthermore, we repeated our analysis on 4-month-old organoids and observed only a small reduction of total *MAPT* mRNA in HSV-1 infected organoids compared to Mock **(Figure 4C)**. Although our 2-month-old organoids had a low percentage of 4R *MAPT* mRNA as expected, in the four-month-old organoids we observed a marked increase in 4R *MAPT* mRNA expression **(Figure 4D)**. Analysis of 4R *MAPT* mRNA showed that HSV-1 infected organoids had a significant increase in the percentage of total *MAPT* mRNA that was 4R *MAPT* when compared to Mock **(Figure 4D).** We next performed immunofluorescence and western blot analyses of 4-month-old organoids infected with either Mock or HSV-1. Immunofluorescence analysis of organoid sections revealed that 4-month-old HSV-1 infected organoids have an increased expression of 4R-Tau protein compared to Mock **(Figure 4E & F)**. It is important to note that it is hard to detect 4R-Tau protein *in vitro* due to low expression in 2D models of iPSC derived neurons therefore further highlighting our observation in a 3D model.^47^ In further support our model, a majority of the 4R-Tau staining in HSV-1 infected organoids was detected at cells that were tdTomato positive indicating a strong link between HSV-1 infection and upregulation of 4R-Tau expression **(Figure 4E)**. We next measured total Tau and 4R-Tau expression levels in organoid protein lysates by western blotting and observed a statistically significant increase in 4R-Tau protein expression in HSV-1 infected organoids compared to Mock **(Figure 4G and H).** This suggests that HSV-1 infection induces both a shift in alternative splicing leading to increased exon 10 containing (4R) *MAPT* mRNA and 4R-Tau protein expression and hyperphosphorylation of Tau protein.

**Figure 4.**
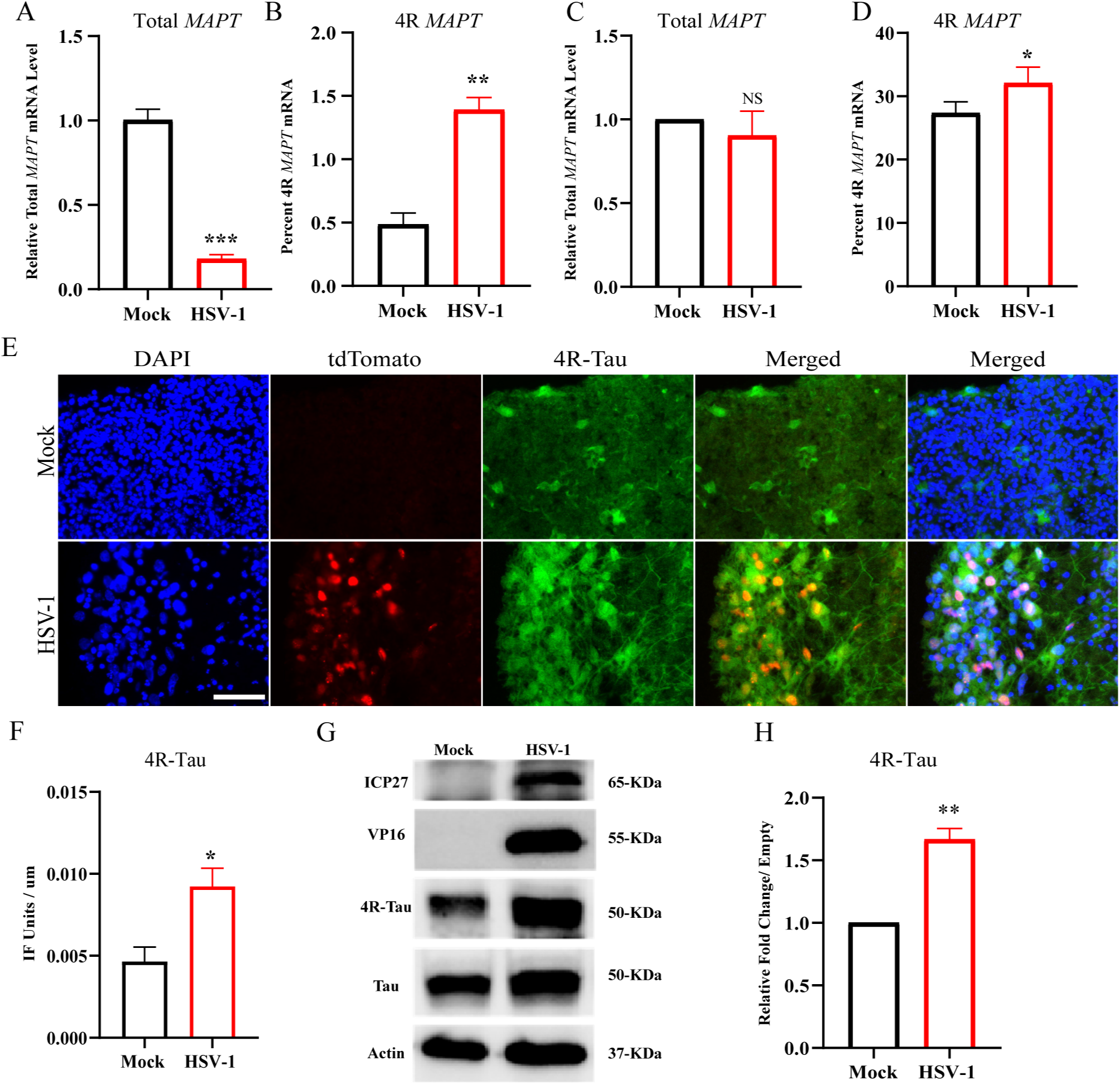
HSV-1 Infection Increases 4R *MAPT* Splicing and 4R-Tau Expression in Forebrain Organoids. A. Reverse Transcription-Polymerase Chain Reaction (RT-PCR) analysis of total *MAPT* mRNA relative to the housekeeping gene hypoxanthine phosphoribosyl transferase 1 (*HPRT1*) in 2-month-old organoids. N of 3, unpaired Student t test. B. Percent 4R *MAPT* mRNA relative to total *MAPT* mRNA. N of 3, paired Student t test. C. RT-PCR analysis of total *MAPT* mRNA relative to the housekeeping gene *HPRT1* in 4-month-old organoids infected. N of 6, unpaired Student t test D. Percent 4R *MAPT* mRNA relative to total *MAPT* mRNA. N of 6, Paired, Student t test. E. Immunofluorescence analysis of 4R-Tau on 4-month-old organoid sections infected with either Mock or HSV-1 tagged with tdTomato. F. Quantification of immunofluorescence signal from E. N of 3.G. Western blot analysis of HSV-1 ICP27, HSV-1 VP16, 4R-Tau protein, total Tau and actin expression in Mock and HSV-1 infected 4-month-old organoids. H. Densitometry quantification of the amount of 4R-Tau relative to Total Tau in the results from G. N of 3, unpaired Student t test. NS = P. Value > 0.05. P* ≤ 0.05, P** ≤ 0.01, P*** ≤ 0.001, P**** ≤ 0.0001. Error bars are SEM. Scale bar is 50 um.

### 3.5. ICP27 is both Necessary and Sufficient to Increase 4R *MAPT* mRNA and 4R-Tau in HSV-1 Infected Neurons

Forebrain organoids represent a complex mix of both neural and glial cell types, and recent findings suggest that astrocytes also express Tau, potentially contributing to altered Tau splicing observed in HSV-1 infected organoids.^48^ Therefore, we wanted to determine if HSV-1 infection alone in neurons would alter splicing of *MAPT* mRNA when compared to Mock controls. We utilized human SH-SY5Y neuroblastoma cells that were differentiated for three weeks using the published protocol in Shipley et al. ^35^ **(Figure 5A)**. These differentiated SH-SY5Y-neurons have extended neurites and are also positive for the neural marker B-Tubulin III **(Figure 5B)**. Importantly differentiated SH-SY5Y-neurons express both *MAPT* mRNA and protein. Using these differentiated cells we compared the effect of HSV-1 infection on *MAPT* exon 10 inclusion. We either Mock infected or infected differentiated SH-SY5Y-neurons with HSV-1 for 24 hours and processed the mRNA and protein for *MAPT* mRNA and protein analysis. RT-PCR analysis revealed that total *MAPT* mRNA was decreased in HSV-1 SH-SY5Y-neurons when compared to Mock controls as expected due to the expression of HSV-1 VHS. This result is very similar to what we observed in the HSV-1 infected organoids where we observed a reduction in total *MAPT* mRNA **(Figure 5B)**. Analysis of 4R *MAPT* mRNA expression revealed a significant increase in the percentage of 4R *MAPT* mRNA over total *MAPT* mRNA in HSV-1 infected SH-SY5Y-neurons compared to Mock **(Figure 5C**, **D and E)**. Western blot analysis further demonstrated a statistically significant increase in 4R-Tau protein expression in HSV-1 infected SH-SY5Y-neurons compared to Mock controls **(Figure 5F and G)**. HSV-1 ICP27 protein is known to interact with proteins involved in alternative splicing of host genes,^23,49^ suggesting a potential role in altering *MAPT* splicing during infection. To test if ICP27 is necessary for the increase in 4R *MAPT*, we infected SH-SY5Y-neurons with either Mock, HSV-1, or HSV-1 lacking ICP27 (Δ-ICP27 HSV-1) for 24 hours. We collected the total cell RNA and protein lysate for further *MAPT* mRNA and protein expression analysis. RT-PCR analysis showed that total *MAPT* mRNA expression levels, which are reduced during HSV-1 infection, is restored to similar levels as Mock in Δ-ICP27 HSV-1 infected SH-SY5Y-neurons **(Figure 6A).** In agreement with Figure 5, we observed that HSV-1 infection increased the total percentage of *MAPT* mRNA that was 4R *MAPT* **(Figure 6A**, **B and C**). However, in the absence of ICP27 we observed that there was no difference in 4R *MAPT* mRNA levels when compared to Mock **(Figure 6A**, **B and C)**. Next, we performed western blot analysis of 4R-Tau and found a significant increase of 4R-Tau protein levels during HSV-1 infection, which was not observed with the Δ-ICP27 HSV-1 virus **(Figure 6D and E)**. These results indicated that ICP27 is necessary, at least in part, for inducing altered splicing of *MAPT* during HSV-1 infection.

**Figure 5.**
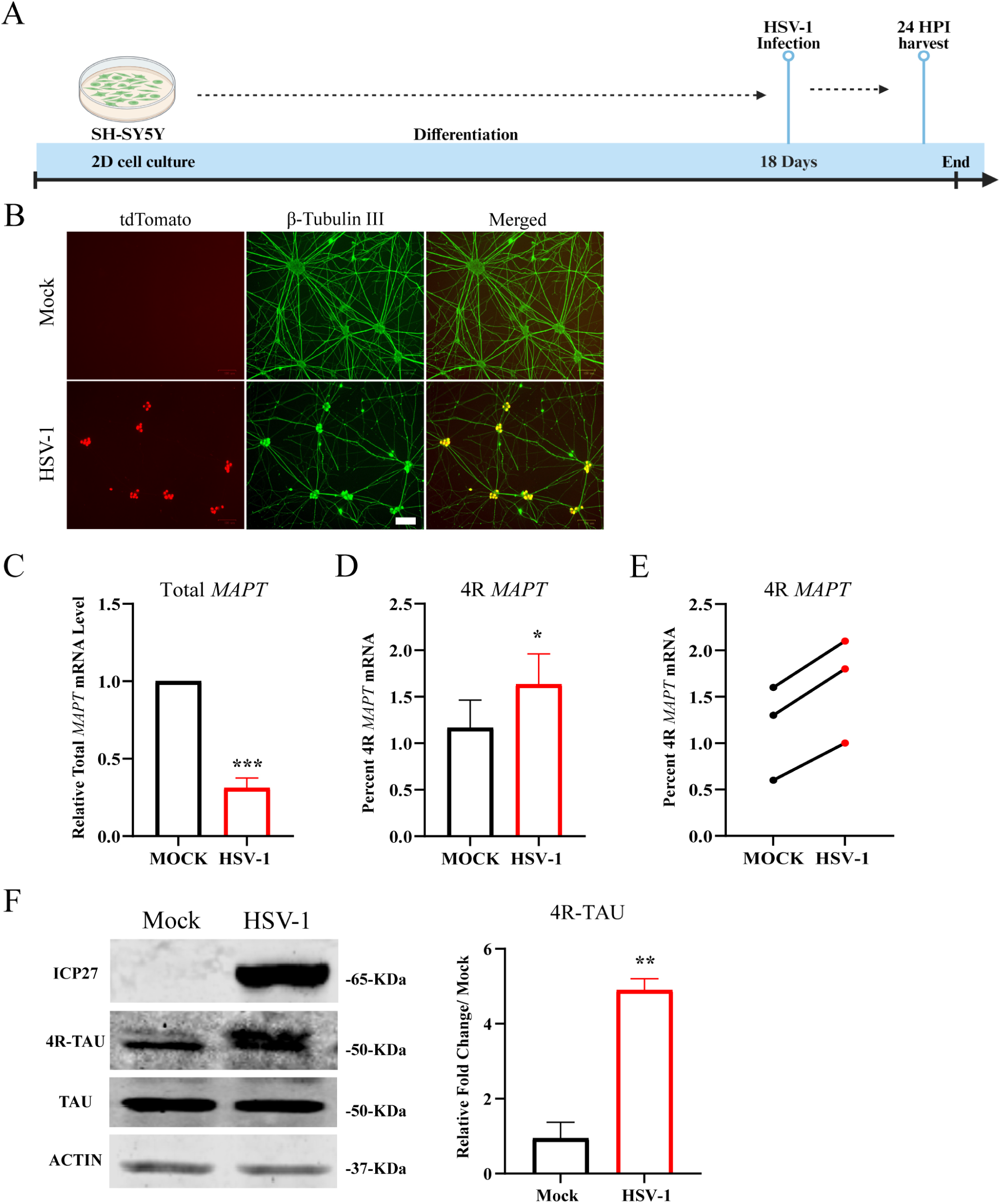
HSV-1 Infection Increases *MAPT* Splicing in 2D Differentiated SH-SY5Y-Neurons. A. Schematic diagram of SH-SY5Y-neurons differentiation and timeline for HSV-1 infection. B. Immunofluorescence analysis of β-Tubulin III in Mock and HSV-1-infected SH-SY5Y-neurons. C. RT-PCR analysis of total *MAPT* mRNA relative to the housekeeping gene *HPRT1*. D. Percent 4R *MAPT* mRNA relative to total *MAPT* mRNA. Paired, Student t test E. Line graph of individual experiments analyzed in D. F. Western blot analysis of HSV-1 ICP27, 4R-Tau protein, total Tau and actin expression in Mock and HSV-1 infected SH-SY5Y-neurons. G. Densitometry quantification of the amount of 4R-Tau relative to Total Tau in the results from F. Unpaired Student t test. All experiments are representative of N of 3 with NS = P. Value > 0.05. P* ≤ 0.05, P** ≤ 0.01, P*** ≤ 0.001, P**** ≤ 0.0001. Error bars are SEM. Scale bar is 100 um.

**Figure 6.**
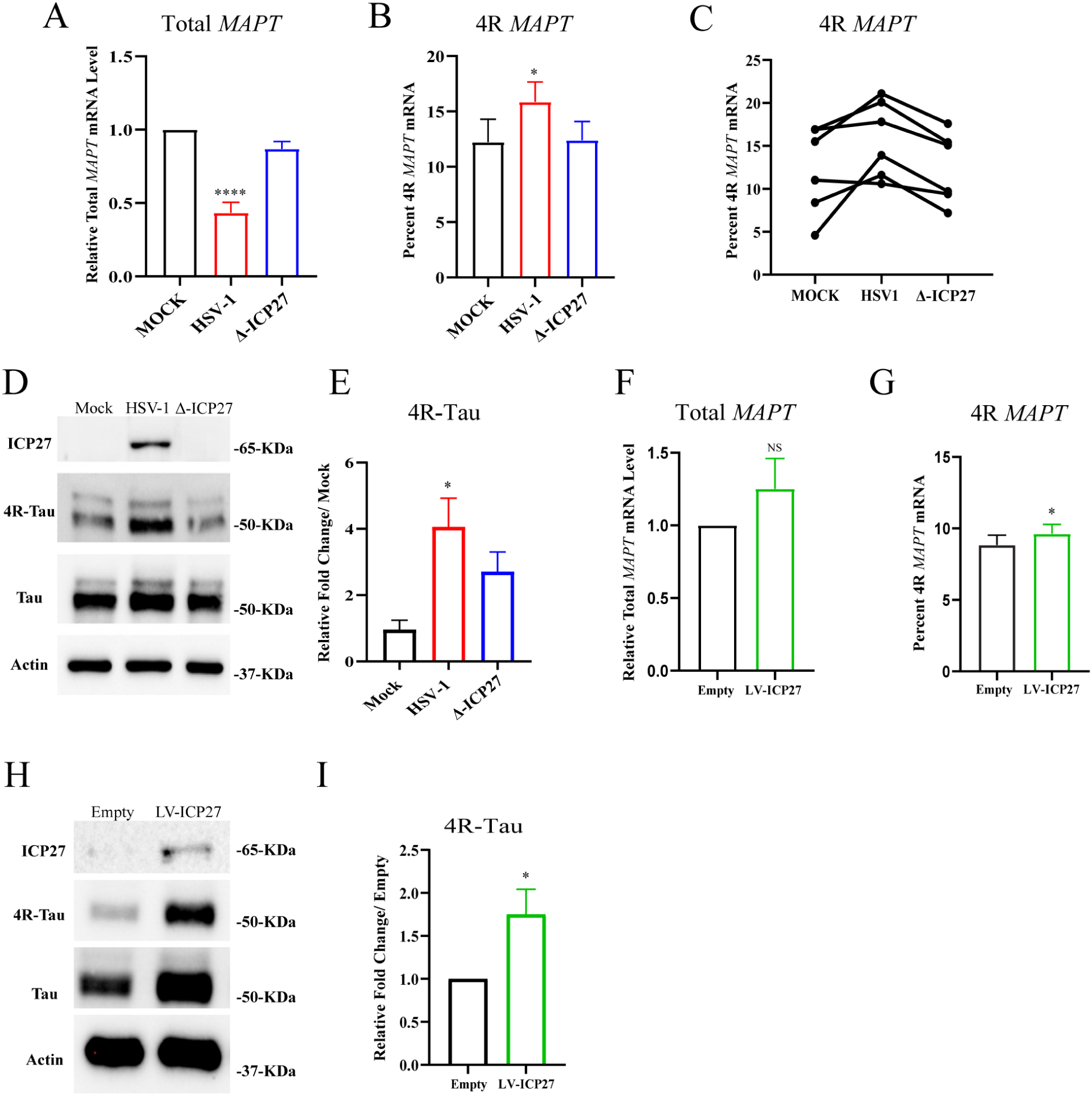
HSV-1 ICP27 Protein is both Necessary and Sufficient to Increase 4R *MAPT* Splicing. A. RT-PCR analysis of total *MAPT* mRNA relative to the housekeeping gene *HPRT1* in Mock, HSV-1 and Δ-ICP27 HSV-1 infected SH-SY5Y-neurons. N of 6. Unpaired Student t test. B. Percent 4R *MAPT* mRNA relative to total *MAPT* mRNA. N of 6. Paired Student t test. C. Line graph of individual experiments analyzed in B. D. Western blot analysis of HSV-1 ICP27, 4R-Tau protein, total Tau and actin expression in Mock, HSV-1 and Δ-ICP27 HSV-1 infected SH-SY5Y-neurons E. Densitometry quantification of 4R-Tau relative to total Tau from the results in D. N of 4. One-way anova. F. RT-PCR analysis of total *MAPT* mRNA relative to the housekeeping gene *HPRT1* in empty vector and ICP27 transduced SH-SY5Y-neurons. N of 4. Unpaired, Student t test. G. Percent 4R *MAPT* mRNA relative to total *MAPT* mRNA. N of 4. Paired Student t test. H. Western blot analysis of HSV-1 ICP27, 4R-Tau protein, total Tau and actin expression in empty vector (Empty) and ICP27 transduced SH-SY5Y-neurons (LV-ICP27). I. Densitometry quantification of 4R-Tau relative to total Tau from the results in G. N of 5. Unpaired Student t test. NS = P. Value > 0.05. P* ≤ 0.05, P** ≤ 0.01, P*** ≤ 0.001, P**** ≤ 0.0001. Error bars are SEM.

After establishing that HSV-1 ICP27 protein was necessary to induce altered splicing of *MAPT*, we next wanted to determine if it is also sufficient to alter *MAPT* splicing. To address this question, we used a Tet-On lentivirus system. This system requires the reverse tetracycline transcriptional activator (rtTA) to function. The rtTA binds to the tetracycline response element (TRE) in the presence of tetracycline or its derivative doxycycline, driving the expression of the target gene, in this case, ICP27. We constructed a tetracycline-inducible gene construct that constitutively expresses GFP and induces ICP27 expression only when doxycycline is present. SH-SY5Y-neurons were transduced with either the rtTA construct alone (empty vector) or with the rtTA and the ICP27 gene construct (LV-ICP27). After twenty-four hours post-transduction, we added doxycycline daily for three days to induce ICP27 expression and then collected samples for RNA and protein analysis. Expression of ICP27 was confirmed by western blot **Figure 6H).** RT-PCR analysis of total *MAPT* mRNA showed that there was no significant difference in overall *MAPT* mRNA in SH-SY5Y-neuron transduced with ICP27 when compared to those transduced with the empty vector **(Figure 6 F)**. This is expected as the transduced cells do not express HSV-1 VHS thus host mRNAs are not inhibited. Importantly, analysis of *MAPT* mRNA showed an increase in 4R *MAPT* mRNA expression over total *MAPT* mRNA in ICP27 transduced SH-SY5Y-neurons compared to those transduced with the empty vector **(Figure 6 G)**. Western blot analysis of 4R-Tau and total Tau protein further revealed that ICP27 transduced SH-SY5Y-neurons had a significant increase in 4R-Tau protein levels relative to total Tau protein levels when compared to SH-SY5Y-neurons transduced with the empty vector alone **(Figure 6H and I).** These results indicate that ICP27 expression is sufficient to induce an increase in 4R *MAPT* splicing and 4R-Tau protein expression in 2D SH-SY5Y-neurons.

### 3.6. HSV-1 Infection Induces Tau Oligomerization

Recent research on AD has highlighted Tau oligomers, which appear before the formation of neurofibrillary tangles, as a major cause of neurotoxicity during tauopathies.^50,51^ In our study, we investigated whether HSV-1 infection could induce Tau oligomerization. Western blot analysis of total Tau protein expression in HSV-1 infected SH-SY5Y-neurons revealed a significant increase in Tau oligomers compared to both Mock-infected and UV-inactivated HSV-1 control groups **(Figure 7A and B)**. The use of UV-inactivated HSV-1 as a control demonstrates that the observed increase in Tau oligomers is specific to conditions that require active viral transcription. To extend these findings beyond 2D neurons, we examined whether a similar effect occurred in a more complex 3D forebrain organoid model. Immunofluorescence analysis using the T22 oligomeric Tau marker showed a significant increase in oligomeric Tau in HSV-1 infected organoids compared to Mock-infected organoids **(Figure 7C and D)**. This suggests that HSV-1 infection can enhance Tau oligomerization in a 3D neural environment, potentially reflecting a more physiologically relevant human brain model. These results from both 2D and 3D neural models show that HSV-1 infection can contribute to increasing Tau oligomerization. Given that Tau oligomers have been considered to be the most pathological form of Tau in a diseased neuron,^51^ our findings strengthen the link between HSV-1 infection and the progression of Tau related diseases such as AD.

**Figure 7.**
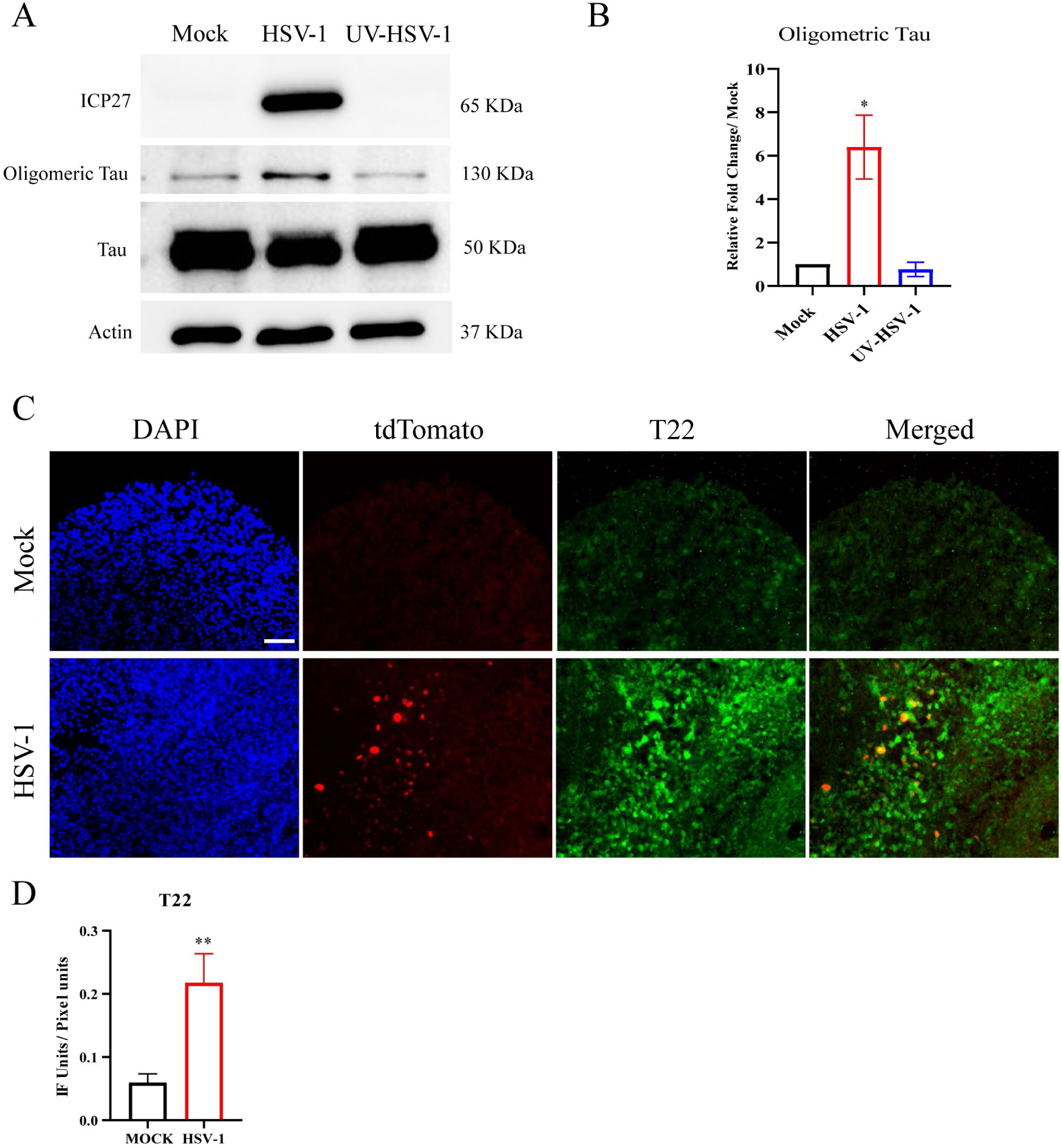
HSV-1 Infection Increases Tau Oligomerization. A. Western blot analysis of HSV-1 ICP27, total Tau, and actin expression in Mock, HSV-1 and UV-inactivated HSV-1 infected differentiated SH-SY5Y-neurons. B. Densitometry of individual replicates from A. N of 3. C. Immunofluorescence analysis of oligomeric Tau marker T22 in Mock and HSV-1 infected brain organoids. D. Immunofluorescence intensity quantification per pixel area for images in C. N of 3. Statistical analysis for all graphs used unpaired Student t test. NS = P. Value > 0.05. P* ≤ 0.05, P** ≤ 0.01, P*** ≤ 0.001, P**** ≤ 0.0001. Error bars are SEM. Scale bar is 50 um.

## 4. Discussion

AD represents the rising cause of dementia and cognitive decline in humans above the age of 65.^52^ With no direct link to any known etiological agent and only approximately five percent of cases attributed to specific mutations to genes (i.e. PSEN1, PSEN2, and APP), there is an immense need to identify the potential disease risk factors.^53^ Neurotropic viruses like HSV-1 have been increasingly implicated in AD progression due to their ability to infect neurons and glial cells and their role in inducing Aβ accumulation. Recently 3D brain organoids have been utilized to model human brains *in vitro.*^27,54^ These organoids share multiple structural and cell type similarities with the human brain and, as such, are increasingly being used to study neural development and neurological diseases like AD.^54^ This model allows investigation into whether HSV-1 can induce changes associated with AD progression, such as Tau hyperphosphorylation and aggregation, in a human-like context. In this study we show that HSV-1 infection leads to an increase in Tau phosphorylation, enhanced 4R *MAPT* splicing, and an accumulation of both 4R-Tau and oligomeric Tau. Our data further indicates that the viral protein ICP27, which is essential for the viral life cycle to be completed, is both necessary and sufficient to induce an increase in both 4R *MAPT* mRNA and 4R-Tau protein expression.

Human forebrain organoids are populated with both neural and glial cell types as early as 2 months. When cultured for 4 months, these organoids typically develop multiple terminally differentiated cell types, including major cortical deep and upper layer neurons, subsets of inhibitory neurons, astrocytes, and oligodendrocyte progenitor cells. Our results show that our 2-month-old organoids had both neural and glial cell types as well as progenitor cells. After infecting these 2-month-old organoids with a GFP-tagged HSV-1, we observed positive GFP signal in cells that also stained positive with NeuN (a neural cell marker), GFAP (a glial cell marker), and Nestin (a neural progenitor cell marker). Further supporting these findings, single-cell RNA sequencing analysis of HSV-1 infected 4-month-old organoids revealed the presence of viral transcripts in neurons, astrocytes, and neuroepithelial cells within our organoids. This gives us further insight into potential cells in the brain that may be contributing towards AD progression as a consequence of HSV-1 infection. Astrocytes, although glial cells, express *MAPT* mRNA and make Tau protein.^48^ Given the close interaction between neurons and astrocytes in the brain our results raise the concern of the possibility that HSV-1 infection in astrocytes could contribute to neuropathology to neighboring neurons.

Tau, a microtubule-associated protein binds to axonal microtubules through its four microtubule binding domains.^55^ The *MAPT* gene, composed of sixteen exons, undergoes alternative splicing to produce six isoforms, classified as 3R or 4R *MAPT* based on the exclusion or inclusion of exon 10.^55^ The Alternative splicing of *MAPT* is mediated by SRSF1 and its Kinase SRPK1. Previously the interaction between HSV-1 protein ICP27 and SRPK1 during infection has been shown to impair phosphorylation of SRSF1 leading to altered splicing of cellular genes like *PML* and nuclear paraspeckle assembly transcript 1 (*NEAT1*).^23,56^ We hypothesized that HSV-1 through its ICP27 protein could alter the splicing of *MAPT* during infection of the brain. To test this hypothesis, we infected 2-month-old forebrain organoids with HSV-1 and compared them to Mock infected organoids. HSV-1 infected 2-month-old organoids had elevated phosphorylated Tau protein and increased 4R *MAPT* mRNA expression compared to Mock-infected controls. Next, we cultured organoids for 4 months allowing them to mature and develop higher levels of 4R *MAPT* mRNA and 4R-Tau protein. After HSV-1 infection of these 4-month-old organoids, we observed a significant increase in both 4R *MAPT* mRNA and 4R-Tau protein compared to Mock-infected controls. Immunofluorescence analysis of our HSV-1 infected organoids showed that HSV-1 tdTomato positive cells had increased 4R-Tau staining within the soma and neurites. HSV-1infection has been previously shown to induce Aβ accumulation and Tau hyperphosphorylation. Our new results show that it also alters splicing of *MAPT* and increases 4R-Tau protein levels.^19,43^ A previous study showed that overexpression of human 4R-Tau in humanized mice resulted in elevated Tau hyperphosphorylation, Tau oligomerization and severe seizures.^57^ This alludes to the possibility that altered *MAPT* expression leading to increased 4R *MAPT* expression over 3R *MAPT* could be an initial step that contributes to increased Tau hyperphosphorylation and oligomerization.

To further explore the effects of HSV-1 infection on *MAPT* splicing in a simplified neural system, we utilized a recently published protocol for differentiating human SH-SY5Y cells into neurons.^58^ Unlike iPSC-derived neurons, which express low levels of 4R-Tau,^59^ human SH-SY5Y cells differentiated into neurons with the addition of retinoic acid (RA), brain-derived neurotrophic factor (BDNF), and nerve growth factor (NGF) exhibit enhanced axonal outgrowth and total Tau expression levels that are comparable to those of the human brain.^60^ This differentiation results primarily in a mixture of cholinergic and dopaminergic neuron-like cells, which are relevant for studying neurodegenerative diseases.^60^ Importantly, *MAPT* splicing in these neurons produces the six major Tau isoforms observed in the mature human brain.^60^ SH-SY5Y-neurons were able to recapitulate our previous findings, exhibiting a significant increase in both 4R *MAPT* mRNA and 4R-Tau protein levels upon HSV-1 infection compared to Mock controls. In this model, HSV-1 infection of differentiated SH-SY5Y-neurons led to a significant increase of 4R *MAPT* mRNA and protein levels relative to Mock-infected controls effectively recapitulating our findings in 3D organoids. This confirmed that HSV-1 infection alone is sufficient to induce increased 4R *MAPT* splicing in neurons. The significance of these results lies in the alteration of the 3R:4R Tau ratio, a feature characteristic of tauopathies. Healthy adult human brains express equal levels of 3R and 4R-Tau isoforms.^61–63^ Recent studies have shown that alteration in the 4R:3R Tau ratio as a result of altered alternative splicing of *MAPT* exon 10, is a key feature of major tauopathies.^55,64,65^ Identification of HSV-1 as a possible etiological factor capable of inducing this alteration in the isotype levels could lead to the discovery of new drug targets and druggable cellular mechanisms responsible for the progression of AD and related tauopathies.

To understand the molecular mechanisms behind the increase in *MAPT* exon 10 splicing and 4R-Tau expression observed HSV-1 infected forebrain organoids and differentiated SH-SY5Y-neurons, we investigated the role of viral protein ICP27. Previous studies have shown that ICP27 interacts with the alternative splicing machinery to regulate host immune protein PML.^23^ Knowing this we hypothesized that ICP27 during HSV-1 infection would alter splicing of *MAPT* exon 10 and lead to an increase of 4R-Tau relative to total Tau. Our results indicated that in the absence of ICP27, HSV-1 infection failed to significantly alter *MAPT* exon 10 splicing, indicating the necessity of ICP27 in inducing the increase in *MAPT* exon 10 inclusion and 4R-Tau expression. To further investigate, we examined whether the expression of ICP27 alone is sufficient to promote increased 4R *MAPT* mRNA expression. We observed that in 2D SH-SY5Y-neurons, expression of ICP27 increases 4R *MAPT* mRNA and 4R-Tau protein expression. These findings enhance our understanding of ICP27’s role in manipulating the host alternative splicing machinery to regulate the splicing of both viral and host gene products such as *MAPT.* This mechanistic insight suggests that ICP27 is a key viral factor contributing to the dysregulation of exon 10 splicing during HSV-1 infection, highlighting its potential as a target for therapeutic intervention in tauopathies.

Additionally, our study reveals that HSV-1 infection increases Tau oligomerization, a process implicated as a driver of Tau pathology.^12,51^ Tau oligomers are neurotoxic species that can propagate between neurons, potentially spreading Tau pathology from a diseased neuron to a healthy neuron. These Tau oligomers are believed to appear before the formation of NFTs.^66^ The ability of HSV-1 to induce the formation of these oligomers suggests a novel mechanism by which viral infection might contribute to the onset and progression of tauopathies like AD. This highlights the potential of HSV-1 as a factor in initiating a cascade of Tau-related neurodegeneration.

In conclusion, we have demonstrated that HSV-1 infection in a 3D forebrain organoid model results in the induction of Tau oligomerization, Tau hyperphosphorylation, and altered splicing of *MAPT.* These results add to our knowledge of what may be the etiological causes of sporadic tauopathies like AD.

## Author Contributions

This study was conceived by Eain Murphy, David Butler and Emmanuel C Ijezie. The first draft of the paper was written by Emmanuel Ijezie. Single cell RNA data preparation and analysis was performed by Micheal Miller, Adam Waickman and Celine Hardy. SH-SY5Y-neurons were maintained and differentiated by Natasha Rugenstein and forebrain organoids were sectioned and stained by Lianna D’Brant and Timothy F Czajka. 4R *MAPT* mRNA and 4R-Tau protein data and results from both 3D and 2D neural models were analyzed by Emmanuel Ijezie. ICP27 null virus Δ-ICP27 HSV-1 was generated by Ava Jarvis.

## Acknowledgments

We would like to thank Dr. Stephen Rice (University of Minnesota), Dr. Greg Smith (Northwestern University) and Dr. Ian Mohr (New York University) for their generous donation of HSV-1 viruses. We also thank Dr. Alison Goate (Mount Sinai) and Julia TCW (Boston University) for the APOE3 and APOE4 isogenic iPSCs, and Dr. Celeste Karch (Washington University) for the healthy control iPSCs that was provided through the generous support of the Tau Consortium of the Rainwater Charitable Foundation. We are grateful to the Regenerative Research Foundation and the NeuraCell Core Facility, directed by Steven Lotz and Dr. Taylor Bertucci of the Neural Stem Cell Institute, for producing the iPSC-derived organoids. This work was funded by the NIH/NIA grant R01AG076007.

## Consent Statement

Consent was not necessary.

