## Supplementary material for "HSV-1 Infection Alters *MAPT* Splicing and Promotes Tau Pathology in Neural Models of Alzheimer’s Disease": Table 1

|  | WESTERN BLOT |  |  |  | WESTERN BLOT |  |  |
| --- | --- | --- | --- | --- | --- | --- | --- |
|  | Primary AB | Catalogue Number | Supplier |  | Secondary AB | Catalogue Number | Supplier |
|  | ICP27 | P1113 | Virusys |  | Anti-Mouse IgG, HRP-linked Antibody | 31430 | Cell Signalling Tecnology |
|  | Tau 4R | 30328S | Cell Signalling Tecnology |  | Anti-Rabbit IgG, HRP-linked Antibody | 7074S | Cell Signalling Tecnology |
|  | Total Tau | MA5-41098 | Thermofisher |  |  |  |  |
|  | VP16 | sc-7545 | Santa Cruz |  |  |  |  |
|  | B-Actin | 4967S | Cell Signalling Tecnology |  |  |  |  |
|  | IFA |  |  |  | IFA |  |  |
|  | Primary AB | Catalogue Number | Supplier |  | Secondary AB | Catalogue Number | Supplier |
|  | Tau | MA5-41098 | Thermofisher |  | Anti-mouse IgG (H+L), F(ab')2 Fragment (Alexa Fluor® 488 Conjugate) | 4408S | Cell Signalling Tecnology |
|  | AT8 | MN1020 | Thermofisher |  | Anti-rabbit IgG (H+L), F(ab')2 Fragment (Alexa Fluor® 488 Conjugate) | 4412S | Cell Signalling Tecnology |
|  | Tau 4R | ET3 | Peter Davies |  |  |  |  |
|  | PThr231 | RZ3 | Peter Davies |  |  |  |  |
|  | MAP2 | 13-1500 | Thermofisher |  |  |  |  |
|  | NEUN | MAB377 | Millipore Sigma |  |  |  |  |
|  | Nestin | MAB1259 | R&D |  |  |  |  |
|  | GFAP | GFAP | Aves Lab |  |  |  |  |
|  | T22 | ABN454 | Millipore Sigma |  |  |  |  |
|  | b-Tubulin III | MA1-118 | Thermofisher |  |  |  |  |

Table 1. List of Primary and Secondary Antibodies.
